# THE HIV-1 SILENCER BCL11B/CTIP2 REGULATES THE TLR3-MEDIATED CELLULAR RESPONSE TO VIRAL INFECTIONS

**DOI:** 10.64898/2026.01.13.699245

**Authors:** Muhammad Kashif, Marco De Rovere, Solene Fenninger, Clementine Wallet, Christian Schwartz, Olivier Rohr, Thomas Loustau

## Abstract

HIV-1 latency remains a major barrier to viral eradication. The persistence of the latently infected reservoirs results from multifactorial mechanisms controlling gene transcription and cellular responses to viral reactivation. The transcriptional repressor BCL11B/CTIP2 is known to promote the establishment and persistence of the latently infected reservoirs. However, its contribution to the cellular response to viral infection has not been investigated before. Together with HEXIM1, paraspeckles contribute to cellular innate sensing of viral infections and to the production of type I interferons. Paraspeckles are nuclear membrane-less organelles composed of proteins, including SFPQ/PSF and NONO, wrapped around the long non-coding RNA NEAT1. Here, we show that HIV-1 infection promotes a TLR3-mediated CTIP2 overexpression delayed from a transitory NEAT1 overexpression, paraspeckles formation and type I IFN production. In response to this interferon production, CTIP2 is recruited to the NEAT1 promotor together with HEXIM1 and repressive histone marks to inhibit NEAT1 gene expression. The resulting depletion of the paraspeckles abrogated INF production, dampening TLR3 signalling and reinforcing HIV-1 latency. Our results demonstrate CTIP2 as a dual-function regulator that silences HIV-1 transcription and restricts antiviral innate immune response as part of a negative feedback loop limiting the interferon response to viral infection.

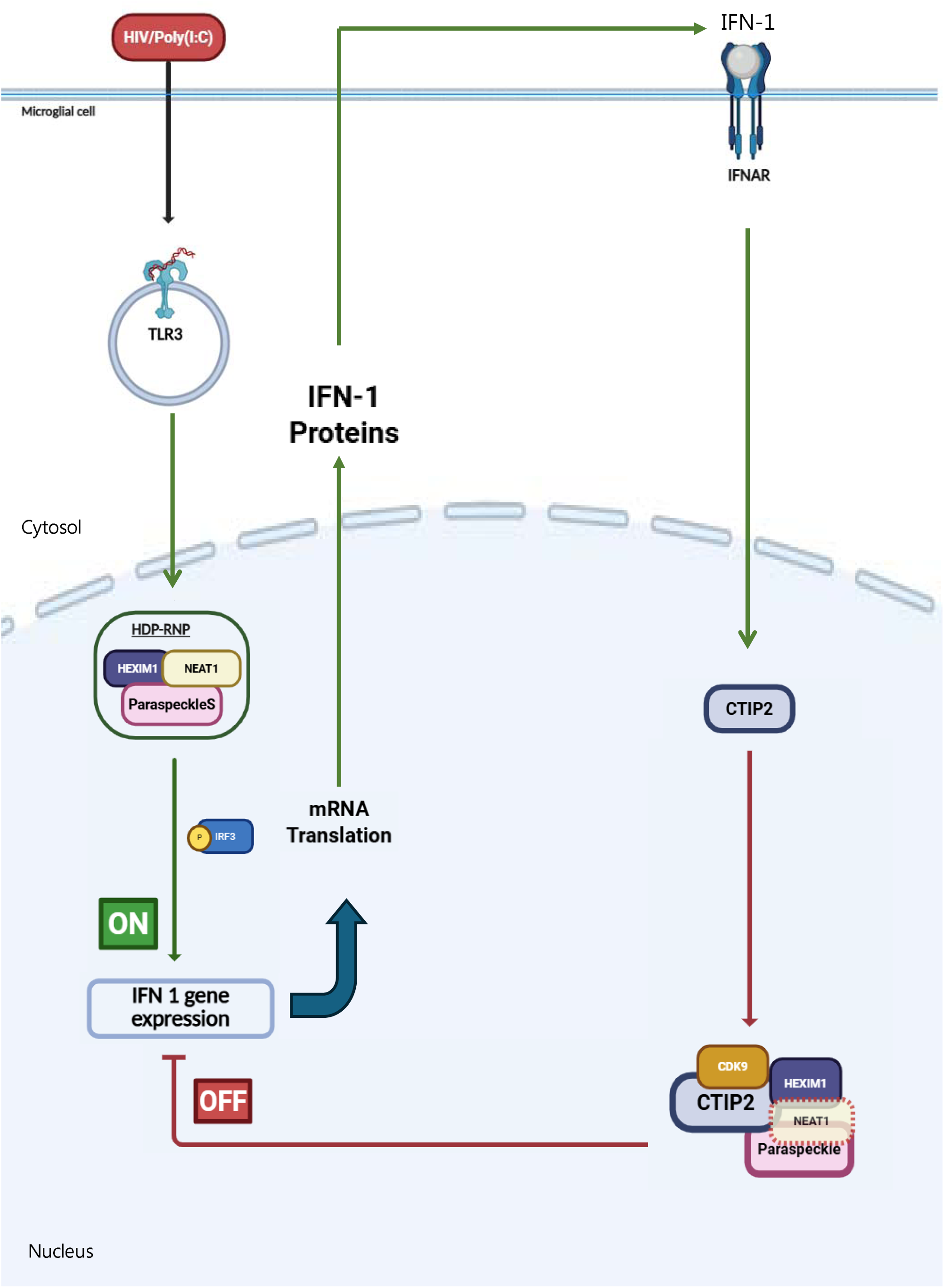

## INTRODUCTION

Despite the success of combination antiretroviral therapy (cART), complete eradication of Human Immunodeficiency Virus type 1 (HIV-1) remains elusive, primarily due to the establishment of latent reservoirs in specific anatomical and cellular compartments ^1–3^. Among these, the central nervous system (CNS) constitutes a unique sanctuary, harboring long-lived cells such as microglia that support viral persistence despite effective systemic viral suppression ^4,5^. Protected by the blood-brain barrier (BBB) and characterized by a tightly regulated immune environment, microglia provide a favorable niche for transcriptionally silent proviral genomes, complicating efforts toward functional or sterilizing cure ^4,6,7^. HIV-1 latency in microglia is maintained through multifaceted mechanisms, including the action of host-derived transcriptional repressors that reshape the epigenetic landscape at the proviral long terminal repeat (LTR). Among these, the transcription factor CTIP2 (also known as BCL11B) has emerged as a pivotal silencer of HIV-1 transcription in CNS-resident cells ^8,9^. CTIP2 exerts its repressive activity by recruiting chromatin-modifying enzymes, including histone deacetylases (HDAC1/2), the histone methyltransferase SUV39H1, and heterochromatin protein 1α (HP1α), to the viral promoter, thereby inducing heterochromatin formation and silencing proviral transcription ^1,10^. By interacting with Hexim-1 and 7SK RNA, CTIP2 also sequesters P-TEFb in an inactive state, preventing cellular gene elongation and HIV-1 transcriptional reactivation ^11,12^. Notably, in microglial models of latency, CTIP2 maintains a deeply repressed proviral state that resists reactivation even in the presence of latency-reversing agents ^5,8,12,13^.

Beyond direct transcriptional repression, emerging evidence implicates CTIP2 in the regulation of host gene expression through interactions with nuclear membrane-less organelles (MLO) and noncoding RNAs ^12^. Among known MLOs, paraspeckles are implicated in various aspects of RNA metabolism and immune regulation, including the nuclear retention of double-stranded RNA and the modulation of transcriptional responses to viral infection and interferon signalling ^14–18^. These dynamic ribonucleoprotein condensates are built around the long noncoding RNA NEAT1 and composed of structural proteins such as NONO, PSPC1, and SFPQ^18–20^. Paraspeckles and NEAT1 interact with several immune regulators and have been shown to coordinate the innate immune response to viral nucleic acids via the HEXIM1-DNA-PK-paraspeckle (HDP RNP) complex, which modulates the cGAS STING IRF3 DNA-sensing pathway ^21^. Along with cGAS, Toll-like receptor 3 (TLR3) is a key sensor of viral double-stranded RNA that signals through the TRIF adaptor to activate TBK1, IRF3, and NF-κB, leading to the production of type I interferons and inflammatory cytokines ^22,23^. Microglial cells express high levels of TLR3, and their activation can reverse HIV-1 latency in some cellular models by removing repressive chromatin barriers ^24–26^. However, persistent TLR3 activation in the CNS is associated with neuroinflammation and cognitive dysfunction in HIV-infected individuals. Of note, HIV-1 has evolved multiple strategies to antagonize TLR signalling and maintain immune quiescence ^27^. NEAT1 expression itself has been shown to influence TLR3 activation and intracellular localization, thereby linking nuclear RNA scaffolds to PRR-mediated signalling ^28^. In this study, we investigate the functional relationship between CTIP2, paraspeckle components, and the TLR3 signalling axis during HIV-1 infection. Using a combination of transcriptomic profiling, RNA-protein interaction assays, and immunological stimulation models, we describe how CTIP2 orchestrates viral silencing and dampens innate immune activation. These results reveal a novel dual role for CTIP2 as a regulator of both viral latency and cellular antiviral signalling, providing insights into potential therapeutic strategies for reactivating latent viruses.

## Results

### 1. CTIP2 Associates with NEAT1 and Paraspeckle Components

We previously identified a BCL11b/CTIP2 CLIP-seq gene set comprising 740 transcripts in HEK293 and microglial cells, with more than 90% corresponding to protein-coding RNAs^29^. This indicated that CTIP2 associates broadly with mRNAs, potentially influencing their processing, nuclear retention, or stability. Genome-wide mapping of CTIP2-bound RNAs by CLIP-seq revealed strong enrichment of the NEAT1 Lnc RNA in both HEK293 (red) and microglial cells (blue) (Figure 1A). Significant CLIP peaks were concentrated across the 5’ region, common to both short and long isoforms of Neat1, whereas IgG controls displayed negligible signal. Replicate experiments confirmed the reproducibility of these binding patterns, supporting specific recognition of NEAT1 transcripts by CTIP2. To validate the RNA-binding activity of CTIP2 and confirm its association with NEAT1 transcripts, we performed RIP-qPCR in human microglial cells using two set of primers named total NEAT1 and NEAT1_2 targeting respectively the common 5’ region of NEAT1-1 and NEAT1_2 isoforms and the central region of NEAT1_2 (Figure 1SA). While Total NEAT1 was robustly recovered, NEAT1_2 showed less but specific enrichments demonstrating the binding of CTIP2 on both isoforms via their common 5’ end sequence (Figure 1B and S1B). In parallel, RIP with SFPQ antibodies revealed the same level of binding on 5’ end region and the central region of NEAT1, confirming the previously reported specific bindings to the NEAT1_2 isoform (Figure 1B and S1B). The binding of CTIP2 at the 5’ end of NEAT1 may suggest that CTIP2 interacts with the paraspeckles components at the periphery of the MLO. Proteomic analysis further supported the integration of CTIP2 into paraspeckle networks. Quantitative mass spectrometry of Flag-CTIP2 IP complexes in HEK293 cells revealed a tightly interconnected cluster of paraspeckle-associated proteins and RNA-binding factors, including SFPQ, NONO, PSPC1, FUS, MATR3, HNRNPD, and CDC5 (Figure 1C). These proteins are known to regulate innate immune responses, RNA splicing, and paraspeckle integrity ^20, 30, 38^. These interactions were highlighted by STRING network analysis, positioning CTIP2 within a functional ribonucleoprotein module. Interactions of paraspeckles components like NONO, SFPQ, PSCPC1, FUS were confirmed by reciprocal co-immunoprecipitation targeting SFPQ in HEK293 cells overexpressing CTIP2, which validates direct associations of CTIP2 with paraspeckle core proteins (Figure 1D) and co-immunoprecipitation targeting Flag-CTIP2 in HEK293 cells overexpressing Flag-CTIP2 (Figure 1SE). We next examined CTIP2’s spatial localization within paraspeckles. Combined RNA FISH for NEAT1 and immunofluorescence (IF) for SFPQ in HEK293 cells expressing RFP-tagged CTIP2 revealed discrete NEAT1-positive foci overlapping extensively with SFPQ, consistent with paraspeckle localization (Figure 1E). CTIP2 displayed partial but reproducible colocalization with NEAT1 and SFPQ within these nuclear bodies, with higher-resolution imaging showing CTIP2 concentrated at the periphery of NEAT1 foci where paraspeckle proteins typically cluster. These findings were corroborated by supplementary co-IF, where SFPQ was observed in nuclear colocalization with CTIP2 upon CTIP2 overexpression, indicating that CTIP2 not only associates with but also reorganizes paraspeckle components (Figure S1D). Together, these biochemical and imaging data provide strong evidence that CTIP2 is physically integrated into paraspeckle subdomains through direct interactions with NEAT1 RNA and paraspeckle proteins.

**Figure 1.**
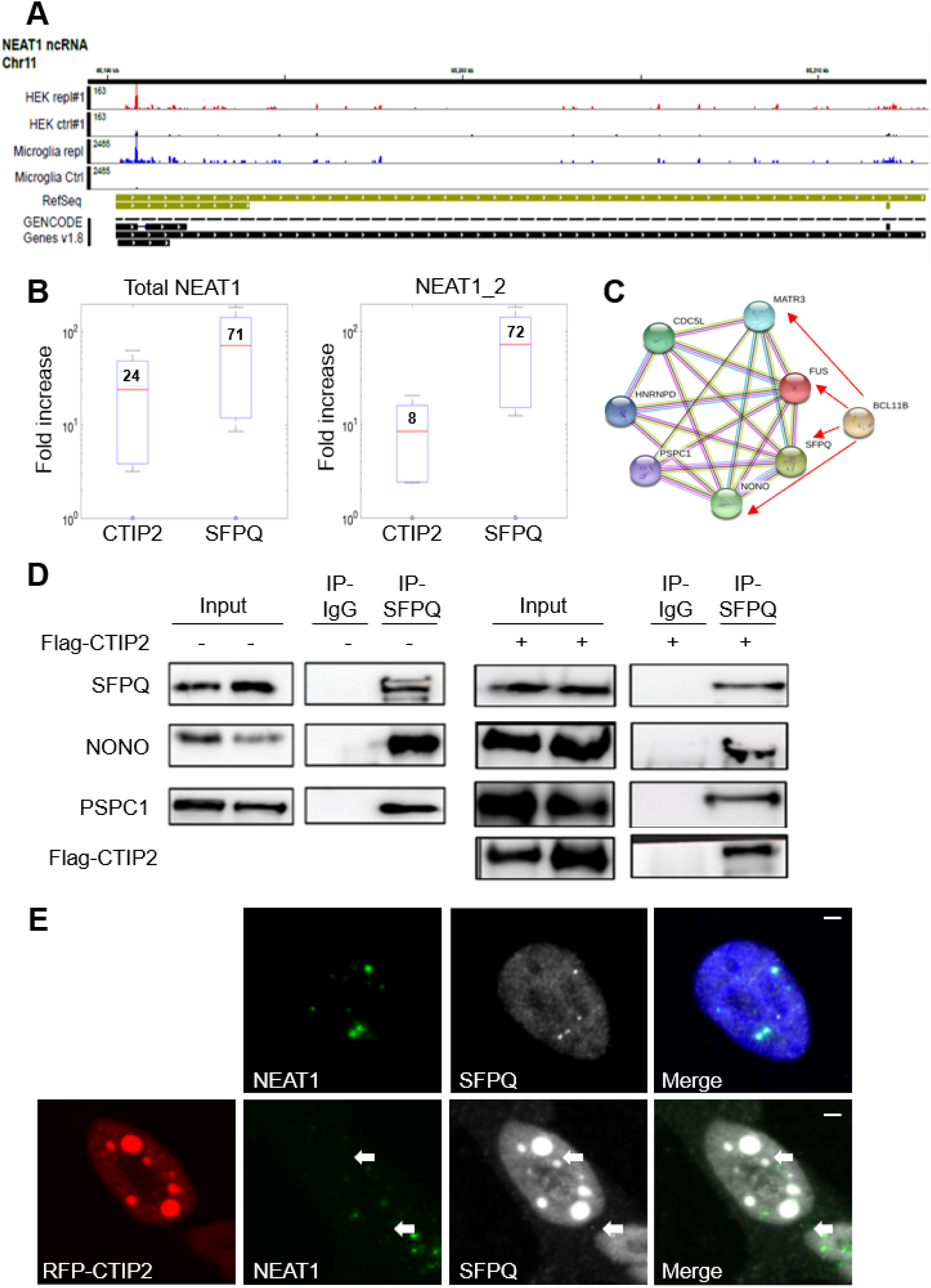
Mapping and Validation of CTIP2-NEAT1 and Paraspeckle Interactions. **(A)** CLIP-seq analysis of CTIP2 and RNA interactions in HEK293 (red) and microglial (blue) cells. Significant binding peaks were detected across the 5′ region of both NEAT1 isoforms compared to IgG controls, indicating specific association of CTIP2 with NEAT1 transcripts. **(B)** RIP-qPCR of microglial cell extracts using CTIP2 and SFPQ antibodies. Both NEAT1_1 and NEAT1_2 isoforms were enriched with CTIP2 since with SFPQ showed preferential enrichment across the central region of NEAT1_2. Binding levels are presented relative to IgG control non-specific bindings. Data represent mean ± SEM from three independent experiments. **(C)** STRING network analysis of Quantitative mass spectrometry analysis Identified a highly interconnected cluster of paraspeckle proteins (SFPQ, NONO, PSPC1) and RNA-binding proteins (FUS, MATR3, HNRNPD, CDC5L) was identified. Proteins indicated by red arrows were selected for further validation. **(D)** Co-immunoprecipitation assays of CTIP2 with the paraspeckles proteins. IP using anti-SFPQ antibodies pulled down CTIP2 and additional paraspeckle components, validating stable interactions of CTIP2 with SFPQ together with NONO and PSPC1. (E) RNA FISH and immunofluorescence in cells expressing RFP-tagged CTIP2. NEAT1 (green) and SFPQ (grayscale) localized to nuclear paraspeckle foci, with partial colocalization of CTIP2 (red). Nuclei were counterstained with DAPI (blue).

### 2. CTIP2 Modulates NEAT1 expression upon TLR3 activation in Microglial cells

RNA-Fish and IF coupling experiments were also performed in microglial cells to visualize the colocalization between CTIP2 and NEAT1 (Figure S2A). This observation proved very difficult, as most cells expressing CTIP2 showed weak NEAT1 staining. The number of paraspeckles per nucleus was quantified and showed a significant reduction in cells overexpressing CTIP2 (Figure S2B), resulting from a reduction in Neat-1 expression of almost 50% in these same cells (Figure S2C). To investigate how CTIP2 and NEAT1 are regulated during viral infection, human microglial cells were infected with a VSVG pseudotyped pNL4.3Δenv HIV-1. Transcript levels were quantified over 4 days by qRT-PCR using isoform-specific primers for Total NEAT1, NEAT1_2, and CTIP2. HIV-1 infection induced a robust and time-dependent upregulation of both NEAT1 isoforms, peaking at 48h and CTIP2 at 72h (Fig. 2A-B, S2D). CTIP2 protein was also markedly elevated in HIV-infected microglial cells at 72h (Fig. 2C). Co-treatment with a TLR3/dsRNA complex inhibitor (100 µM) abolished both Neat1 and CTIP2 inductions, linking the response to TLR3-mediated viral sensing (Fig. 2A-B, S2D). These results extend prior studies demonstrating that viral dsRNA sensing through TLR3 induces NEAT1 ^30^, while identifying CTIP2 as a co-regulated factor in the same pathway. To mimic TLR3-mediated response to HIV-1 infection under controlled conditions, we transfected microglial cells with synthetic poly(I:C), comparing high molecular weight (HMW) and low molecular weight (LMW) duplexes. To confirm the stimulation of the TRL3-mediated response upon stimulation of microglial cells with Poly(I:C), IRF3 phosphorylation was assessed by Western blot. As expected, a strong and early induction of IRF3 phosphorylation was detected with both LMW and HMW Poly(I:C) (Fig. S2E). HMW poly(I:C) induced a notably stronger expression of Total NEAT1 and NEAT1 long isoform at 48h than LMW poly(I:C), consistent with the previous infection results and higher potency of HMW duplexes in engaging TLR3 ^34^ (Fig. 2D and S2E). Poly(I:C) also induced CTIP2 overexpression, with the highest at 72h post-transfection. Once again, this upregulation is more pronounced with the use of HMW form (Figure 2E). This observation is confirmed at the protein level by Western blot (Fig. 2F) and by ELISA essays showing the kinetics of CTIP2 protein expression in nuclear extracts from microglial cells over time (Fig. 2G). At 48h post Poly(I:C) HMW stimulation, RNA-FISH revealed abundant NEAT1 expression distributed throughout the nucleoplasm in much bigger structures than the unstimulated paraspeckles shown in Figure 1H, while CTIP2 immunostaining appeared diffuse and weak (Fig. 2H). However, at 72h, the CTIP2 signal became more intense and redistributed into characteristic punctate nuclear foci, whereas the NEAT1 signal was largely reduced. Importantly, these CTIP2 puncta do not overlap but remain spatially close to NEAT1 condensates. Taken together, these results show an inverse relationship in the expression of CTIP2 and NEAT1. To define whether CTIP2 modulation actively modifies NEAT1 expression, microglial cells were co-transfected with poly(I:C) and either shRNA constructs targeting CTIP2 (shCTIP2) or a control shRNA (shCtrl). Knockdown efficiency was validated by qRT-PCR and Western blot, which confirmed near-complete depletion of CTIP2 protein (Fig. 2I; Fig. S2G). Under these conditions, poly(I:C)-induced NEAT1 expression was significantly elevated compared to shCtrl cells (Fig. 2J; Fig. S2H). Conversely, overexpression of Flag-tagged CTIP2 (validated by western blotting, Fig. S2I) led to earlier and stronger ectopic CTIP2 gene expression (Figure 2K) but markedly suppressed poly(I:C)-induced NEAT1 upregulation (Fig. 2L; Fig. S2J).

**Figure 2.**
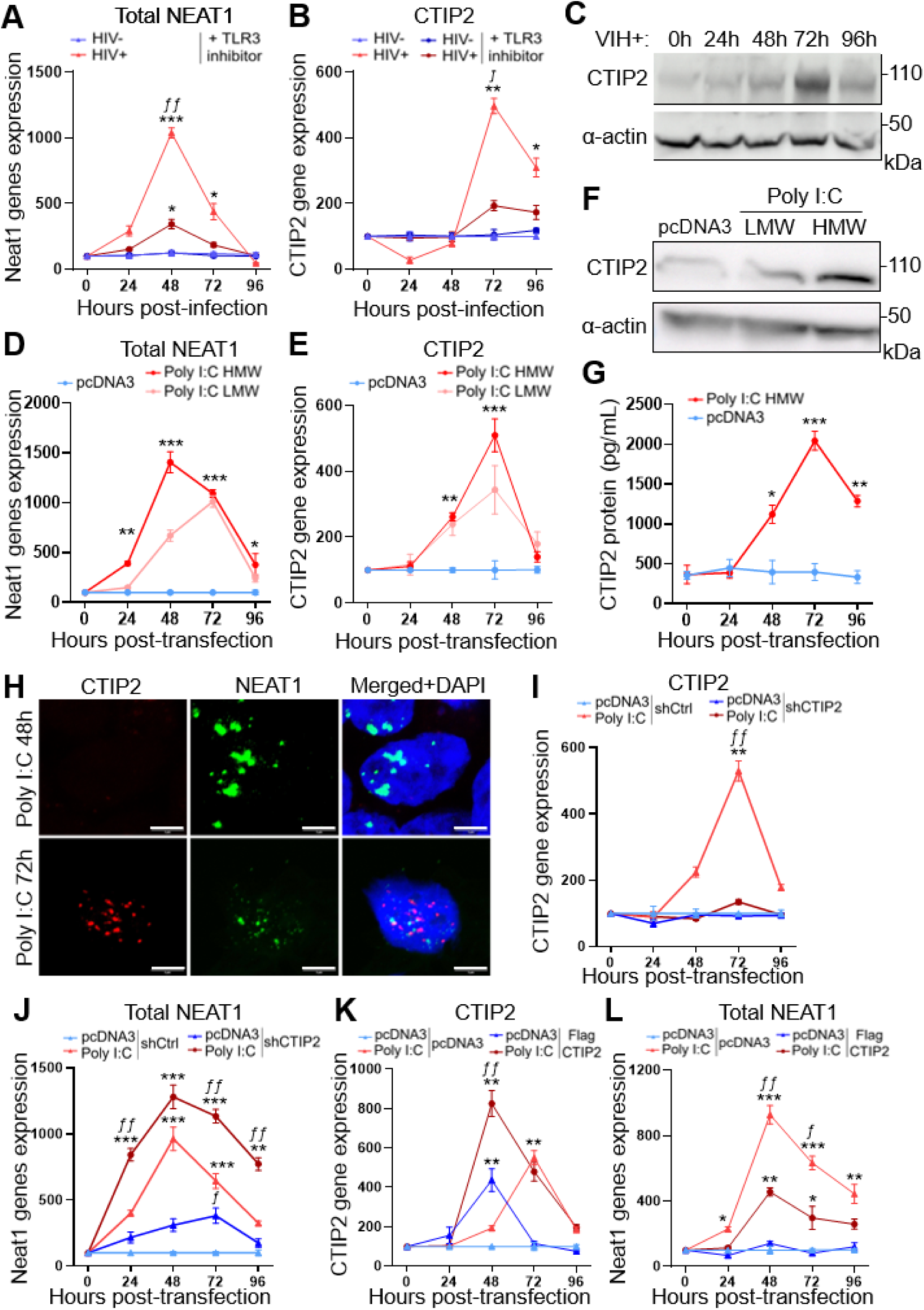
HIV-1 and Poly(I:C) regulate NEAT1 and CTIP2 expression via TLR3 signalling. **(A-B)** qRT-PCR analysis of microglial cells infected with pNL4.3Δenv-VSVG HIV (HIV+) or mock-treated (HIV-), in the presence or absence of a TLR3/dsRNA inhibitor (100 µM). HIV-1 infection induced robust upregulation of Total NEAT1 **(A)** and CTIP2 **(B)** transcripts, peaking at 48-72 h, while TLR3 inhibition abrogated this response. **(C)** Western blot validation of CTIP2 protein levels in HIV-1 infected microglial cells. CTIP2 protein increased at 72 h post-infection, consistent with mRNA data, but was abolished under TLR3 inhibition. α-actin served as a loading control. **(D)** Poly(I:C) stimulation of microglial cells induced strong expression of NEAT1_1, peaking at 48 h. High molecular weight (HMW) poly(I:C) triggered stronger induction than low molecular weight (LMW), while no induction was observed in pcDNA3 controls. **(E-G)** qRT-PCR **(E)**, Western blot **(F)**, and ELISA **(G)** analysis of CTIP2 expression following poly(I:C) stimulation. CTIP2 mRNA and protein levels were significantly upregulated, peaking at 72 h, with HMW poly(I:C) eliciting stronger responses than LMW. Protein kinetics measured by ELISA mirrored transcript induction. **(H)** Confocal microscopy of microglial cells transfected with poly(I:C), showing CTIP2 (red) and NEAT1 RNA foci (green). At 48 h, NEAT1 foci were abundant with weak CTIP2 staining, while at 72 h CTIP2 accumulated in punctate foci overlapping with NEAT1-positive paraspeckles. Nuclei were stained with DAPI (blue). Scale bars, 5 μm. (I-J) qRT-PCR analysis of CTIP2 knockdown in poly(I:C)-stimulated microglia. shCTIP2 efficiently reduced CTIP2 expression **(I)** and resulted in markedly elevated NEAT1 induction at 48 h compared to shCtrl **(J)**. (K-L) qRT-PCR analysis of microglial cells transfected with pcDNA3 or Flag-CTIP2, followed by poly(I:C) stimulation. Overexpression of Flag-CTIP2 led to earlier and higher CTIP2 gene accumulation at 48h (K) but significantly blunted total NEAT1 induction compared to controls **(L)**. Mean ± SEM, n=4 independent experiments per time point and per conditions, multiple Mann-Whitney test. * p < 0.05, ** p < 0.01, *** p < 0.005, **** p < 0.001 (relative to pcDNA3 condition). ƒ pvalue < 0.05, ƒƒ p < 0.01 and ƒƒƒ p < 0.005 (relative to TLR3 inhibitor (A-B), shCtrl **(I-J)** and pcDNA3 (K-L) conditions).

### 3. CTIP2 Stabilizes High Molecular Weight Complexes and Recruits Transcriptional Regulators to the NEAT1 Promoter

We carried out biochemical fractionation of nuclear complexes by glycerol gradient sedimentation and chromatin immunoprecipitation (ChIP-qPCR) targeting the NEAT1 gene promoter to investigate how CTIP2 contributes to nuclear remodeling of protein complexes and transcriptional regulation of NEAT1. For glycerol gradient sedimentation, nuclear extracts from microglial cells transfected with control pcDNA3, with shCtrl and poly(I:C), or with shCTIP2 and poly(I:C) were fractionated on 10-40% glycerol gradients. After ultracentrifugation, 23 fractions were collected from top (lighter) to bottom (heavier) and analyzed by SDS-PAGE and Western blotting. In the control condition, paraspeckle proteins (SFPQ, NONO, PSPC1) and the inactive P-TEFb complex (HEXIM-1, CDK9) were mainly detected in fractions centered on fraction 9, suggesting complexes with comparable sizes. Following 72 h of poly(I:C) stimulation which strongly induces CTIP2 expression, these proteins shifted toward heavier fractions, consistent with the assembly of higher-order nuclear complexes. Importantly, CTIP2 knockdown prevented this redistribution, leaving paraspeckle proteins and transcriptional regulators confined to lighter fractions (Figure 3A). These results suggest that TLR3-mediated induction of CTIP2 reshaped NEAT1-associated factors structures into higher molecular weight complexes. To connect these structural changes to the transcriptional control of NEAT1 gene expression, we performed ChIP-qPCR targeting the NEAT1 gene promoter. At 24 h post poly(I:C) stimulation, we observed increased recruitment of RNA Pol II, p65, and H3Ac at the promoter region, consistent with euchromatin formation, promoter activation, and active NEAT1 transcription (Fig. 3B lower panel). By contrast, at 72 h, coinciding with maximal CTIP2 induction, we detected strong recruitment of CTIP2 and HEXIM-1 (Figure 3B, higher panel), together with increased H3K9 and HDAC1 occupancy, and reduced H3Ac, RNA Pol II and p65 binding. This profile suggests chromatin compaction, P-TEFb inactivation by the recruitment of HEXIM1 together with CDK9 and Cyclin T1, and silencing of the NEAT1 gene promoter, consistent with the above shown qPCR quantifications of NEAT1 expression at 72h post stimulation. Importantly, this poly(I:C) induced dynamic of recruitments was not observed outside the promoter (5′ from the NEAT1 promoter within the XBP1 gene region) (Fig. S3A). Thus, CTIP2 appears specifically at 72 h post-stimulation, when NEAT1 promoter activity is being repressed. Finally, to confirm whether CTIP2 recruitment to the NEAT1 promoter is conserved across cell types, we transfected HEK293T cells with Flag-CTIP2 or pcDNA3 control. ChIP-qPCR confirmed enrichment of CTIP2 and HEXIM-1 at the NEAT1 promoter region in Flag-CTIP2 expressing cells compared to controls (Fig. 3C). No enrichment was detected outside the promoter (Fig. S3B). Our results suggest that CTIP2 expression stimulated by LTR3-mediated sensing promotes the formation of high molecular weight complexes, including paraspeckle-associated complexes, and represses NEAT1 expression by inducing heterochromatin structure at the NEAT1 promoter and inhibiting P-TEFb functions as previously shown for the HIV-1 promoter ^8, 11^.

**Figure 3.**
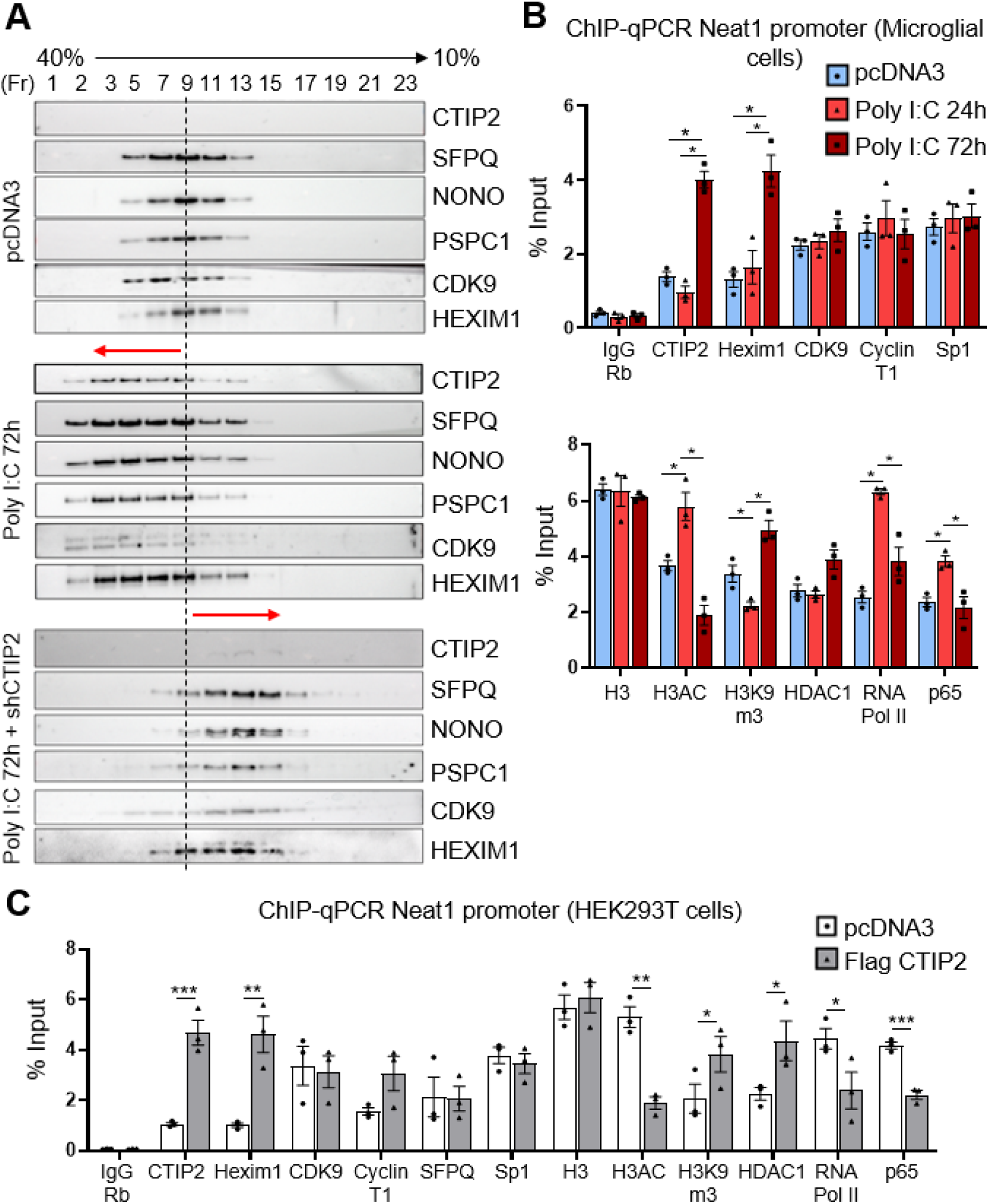
CTIP2 recruits paraspeckle proteins into large complexes and represses NEAT1 promoter activity. **(A)** Glycerol gradient sedimentation of nuclear extracts from microglial cells transfected with pcDNA3 (upper), shCtrl + poly(I:C) (middle), or shCTIP2 + poly(I:C) (lower). Fractions (1-23) were analyzed by SDS-PAGE and Western blotting with the indicated antibodies. Poly(I:C) stimulation shifted paraspeckle proteins and transcriptional regulators toward heavier fractions, consistent with higher-order complex assembly, whereas CTIP2 knockdown abolished this redistribution. Red arrows indicate fraction shifts. **(B)** ChIP-qPCR analysis of microglial cells transfected with pcDNA3 or stimulated with poly(I:C) for 24 h or 72 h. Enrichment of CTIP2, HEXIM1, CDK9, Cyclin T1, and Sp1 (Upper Panel), or HDAC1, RNA Pol II, p65, and histone modifications (H3, H3Ac, H3K9) (Lower Panel), at the NEAT1 promoter region. Data are mean ± SEM from three independent experiments performed in duplicate; significance determined by Mann-Whitney tests (*p ≤ 0.05; **p ≤ 0.01; ***p ≤ 0.001). **(C)** ChIP-qPCR analysis of HEK293T cells transfected with pcDNA3 or Flag-CTIP2 plasmids for 48 h. CTIP2 and HEXIM1 occupancy was quantified at the NEAT1 promoter. Results represent three independent experiments in duplicate, shown as percentage of input. Mean ± SEM, Mann–Whitney tests (*p ≤ 0.05; **p ≤ 0.01; ***p ≤ 0.001).

### 4. CTIP2 Establishes a Negative Feedback Loop to TLR3-Mediated Type I Interferon Responses

Given the importance of paraspeckles and Neat-1 in activating the antiviral immune response and interferon production, measurement of both transcript and protein outputs of IFNβ and IFNα has been performed in microglial cells subjected to HIV-1 infection, Poly(I:C) stimulation, and genetic modulations of CTIP2 expression in microglial cells. Infection of microglial cells with HIV-1 induced a strong but transient increase in IFNβ and IFNα mRNA, peaking at 48 h and declining by 72 h (Figure 4A-B). This induction was abolished by a pharmacological inhibitor of the TLR3/dsRNA complex (100 µM), confirming that HIV-1 engages canonical TLR3 signalling. Parallel analysis revealed that NEAT1 isoforms followed a similar pattern, peaking at 48 h, while CTIP2 mRNA and protein levels rose more slowly, reaching maximal accumulation at 72 h (Figure 2A-C). This temporal separation suggested that CTIP2 induction lags behind interferon production, consistent with a delayed repressive role. To mimic viral sensing, we stimulate microglial cells with high molecular weight (HMW) or low molecular weight (LMW) poly(I:C). Both ligands induced strong IFNβ and IFNα expression, peaking at 48 h, with HMW poly(I:C) eliciting the stronger response (Figure 4C-D). NEAT1 was co-induced in the same time window, while CTIP2 protein accumulation peaked later at 72 h (Figure 2D-F). These kinetics once more suggested a pathway in which CTIP2 upregulation follows TLR3 activation, which first triggers NEAT1 and interferons. We co-transfected poly(I:C) and either shCTIP2 or shCtrl into microglial cells to see if CTIP2 expression affected this pathway. IFNβ and IFNα induction was transient, peaking at 48 h and declining thereafter in shCtrl cells. On the other hand, interferon expression was greatly increased and sustained by CTIP2 knockdown, with increased transcript levels persisting at 72 hours (Figure 4E-F). Conversely, TLR3-mediated reactions were inhibited by Flag-CTIP2 overexpression. Poly(I:C) strongly induced IFNβ and IFNα in control cells, peaking at 48 hours, while by adding CTIP2 overexpression, those inductions are attenuated at all time points (Figure 4G-H). This suppression parallel reduced NEAT1 expression under the same conditions (Figure 2L, S2J), supporting a model where CTIP2 represses interferons indirectly through NEAT1 regulation. Finally, functional assays at the protein level confirmed these effects. ELISA of supernatants collected 72 h post-stimulation showed that CTIP2 knockdown enhanced secretion of IFNβ and IFNα, while CTIP2 overexpression markedly reduced their release (Fig. 4I-J). These results establish CTIP2 as a delayed-acting negative regulator of TLR3-mediated interferon responses, operating through repression of NEAT1 and stabilization of repressive nuclear assemblies.

**Figure 4.**
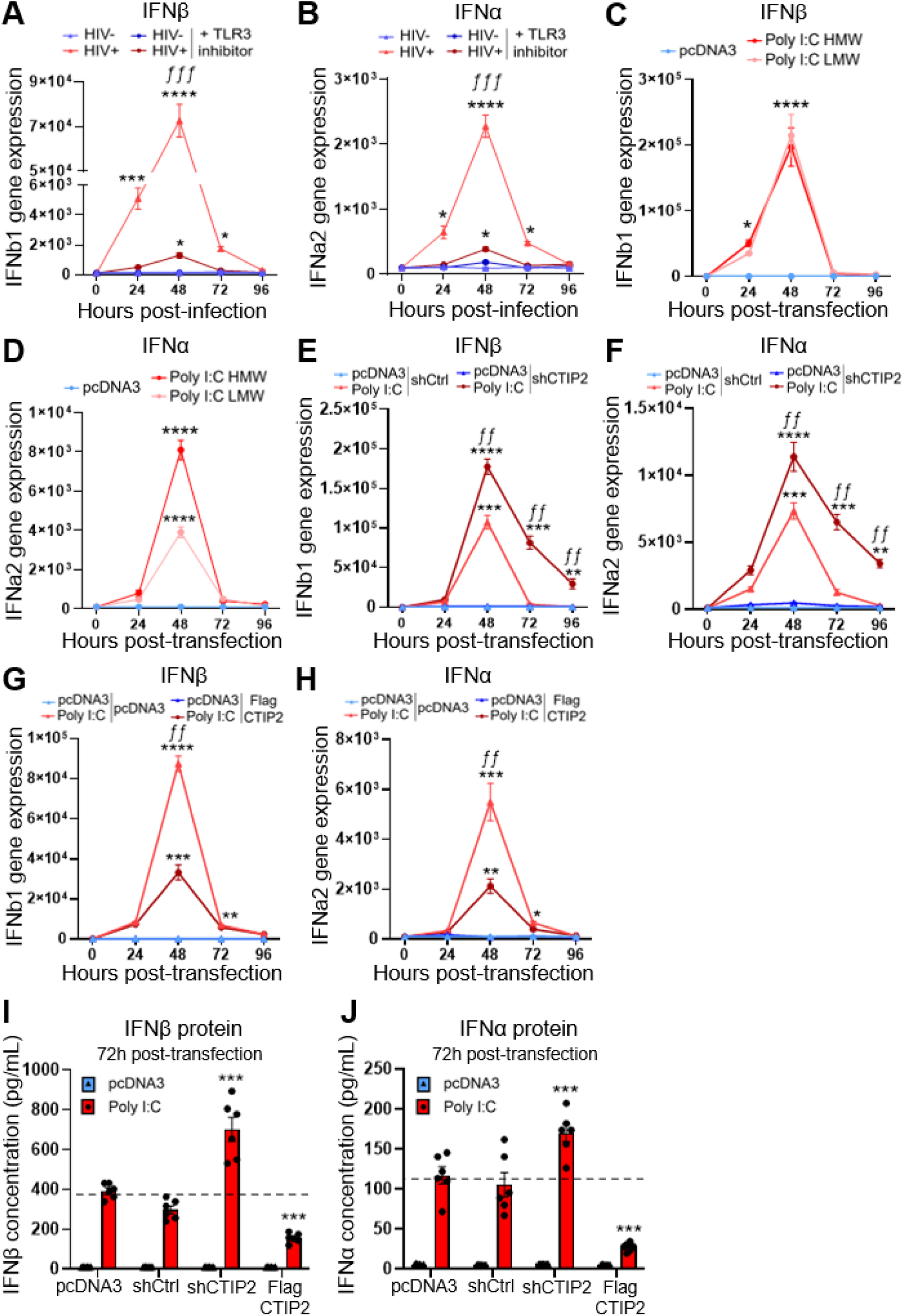
CTIP2 modulates TLR3-dependent type I interferon responses in microglial cells. **(A-B)** qRT-PCR analysis of IFNβ **(A)** and IFNα **(B)** transcripts in microglial cells infected with HIV-1 (HIV+) or mock-treated (HIV-), with or without a TLR3/dsRNA inhibitor (100 µM). HIV-1 induced transient interferon expression peaking at 48 h, which was abolished by TLR3 inhibition. **(C-D)** qRT-PCR analysis of IFNβ **(C)** and IFNα **(D)** in microglial cells transfected with high molecular weight (HMW) or low molecular weight (LMW) poly(I:C). Both isoforms were strongly induced, with HMW eliciting a greater response. **(E-F)** qRT-PCR analysis of IFNβ **(E)** and IFNα **(F)** in microglial cells co-transfected with poly(I:C) and either shCtrl or shCTIP2. CTIP2 depletion enhanced and prolonged interferon expression compared to controls. **(G-H)** qRT-PCR analysis of IFNβ **(G)** and IFNα **(H)** in microglial cells transfected with poly(I:C) and either pcDNA3 or Flag-CTIP2. Overexpression of CTIP2 significantly blunted poly(I:C)-induced interferon expression. **(A-H)** Mean ± SEM, n=4 independent experiments per time point and per conditions, multiple Mann-Whitney test. * p < 0.05, ** p < 0.01, *** p < 0.005, **** p < 0.001 (relative to pcDNA3 condition). ƒƒ p < 0.01 and ƒƒƒ p < 0.005 (relative to TLR3 inhibitor (A-B), shCtrl **(E-F)** and pcDNA3 **(G-H)** conditions). (I-J) ELISA quantification of IFNβ (I) and IFNα **(J)** protein secretion in microglial supernatants 72 h post-transfection. CTIP2 depletion increased, whereas CTIP2 overexpression reduced, interferon protein levels. Data represent mean ± SEM from three independent experiments performed in duplicate, multiple Mann-Whitney test, *** p < 0.005 relative to Poly I:C/pcDNA3 condition (level represented by a dotted line).

### 5. IFNAR Signalling Promotes CTIP2 Induction while Repressing NEAT1 and Type I Interferons

Microglial cells were transfected with HMW poly(I:C) to activate TLR3 and cultured with either IgG isotype control or neutralizing anti-IFNAR1 antibody (100 ng/mL) throughout stimulation. We wanted to test whether type I interferon signalling itself drives CTIP2 expression, since our results showed that CTIP2 induction followed rather than overlapped with NEAT1 and interferon responses during HIV-1 infection and poly(I:C) stimulation. qRT-PCR analysis revealed that poly(I:C) induced strong CTIP2 transcript accumulation in control-IgG treated cells, consistent with our earlier findings. However, IFNAR1/2 blockade reduced CTIP2 induction by more than half, demonstrating that interferon signalling is necessary to sustain poly(I:C)-induced CTIP2 gene expression transcription (Figure 5A). At the protein level, confocal microscopy confirmed these results: poly(I:C)-stimulated cells displayed robust nuclear CTIP2 puncta in the IgG condition, whereas neutralizing IFNAR1/2 markedly reduced both signal intensity and the number of CTIP2-positive foci (Figure 5B). Quantitative image analysis confirmed a significant decrease in CTIP2-positive nuclear area per cell under IFNAR inhibition (Figure 5C). These observations were validated by ELISA, which showed substantially reduced CTIP2 protein levels in microglial cells nuclear extracts upon IFNAR blockade compared to controls (Figure 5D). We next assessed NEAT1 gene expression under these conditions, reasoning that reduced CTIP2 expression would relieve its repression of paraspeckle activity. Indeed, qRT-PCR revealed significant upregulation of both total NEAT1 and its long isoform NEAT1_2 in cells treated with anti-IFNAR1 antibody relative to controls (Figure 5E-F). Thus, IFNAR signalling not only sustains CTIP2 induction but also indirectly limits NEAT1 transcription. Finally, we analyzed type I interferon responses. In IgG-treated controls, poly(I:C) induced robust IFNβ and IFNα expression, consistent with canonical TLR3 signalling (Fig. 4C-D). Importantly, blocking IFNAR1/2 further elevated both IFNβ and IFNα transcript levels (Figs. 5G-H), demonstrating that CTIP2 normally functions as a negative feedback regulator to constrain interferon production. Together, these results establish a self-limiting regulatory circuit in which TLR3 activation initially induces NEAT1 and interferons, which then act through IFNAR signalling to drive CTIP2 expression. Once induced, CTIP2 represses NEAT1 and interferons, thereby dampening antiviral responses. This feedback loop positions CTIP2 as a critical mediator of interferon homeostasis in microglial cells, preventing uncontrolled inflammation while promoting HIV-1 persistence.

**Figure 5.**
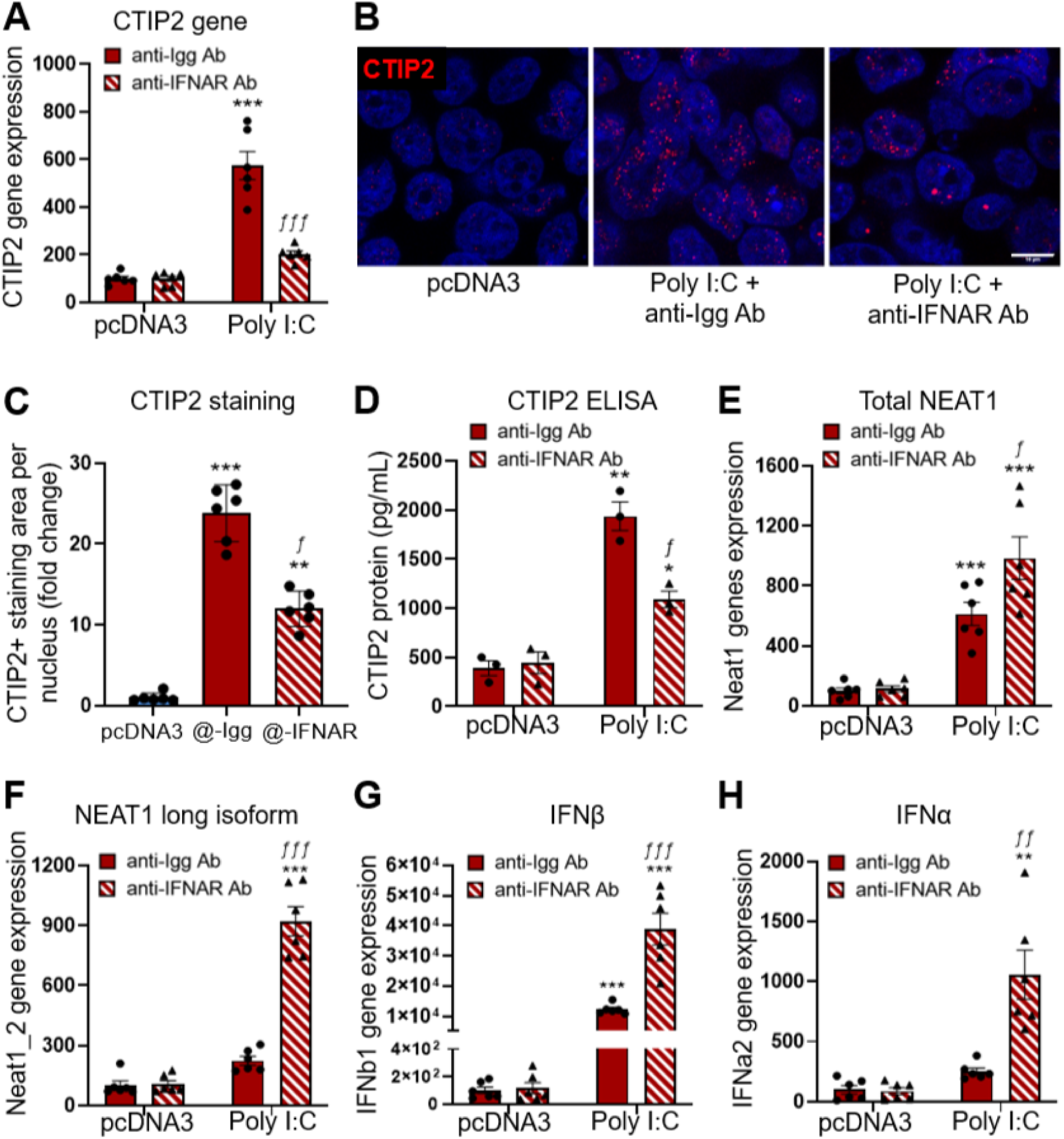
IFNAR Signalling Sustains CTIP2 Induction and Represses NEAT1 and Type I Interferons. **(A-D)** Poly(I:C)-stimulated microglial cells were treated with IgG control or anti-IFNAR1/2 antibody (100 ng/mL). **(A)** qRT-PCR showed that IFNAR blockade reduced CTIP2 mRNA induction by more than half. **(B)** Confocal microscopy of CTIP2 (red) and DAPI (blue) revealed nuclear accumulation in control cells, which was reduced upon IFNAR inhibition. Scale bar, 10 μm. **(C)** Quantification of CTIP2+ nuclear area confirmed reduced nuclear localization. **(D)** ELISA analysis validated reduced CTIP2 protein accumulation under IFNAR blockade. Cells from 5 images per condition have been analyzed. Friedman test, ** p < 0.01, *** p < 0.005 (relative to pcDNA3 condition). ƒ pvalue < 0.05 (relative to anti-IFNAR condition). **(E-F)** qRT-PCR analysis of NEAT1 transcripts in Poly(I:C)-stimulated microglia treated with IgG or anti-IFNAR. IFNAR neutralization increased **(E)** total NEAT1 and **(F)** NEAT1_2 expression compared to controls. **(G-H)** qRT-PCR analysis of IFNβ **(G)** and IFNα (H) in Poly(I:C)-stimulated microglia showed further upregulation under IFNAR blockade, indicating that CTIP2 induction by IFNAR signaling normally restrains interferon output. **(A, D-H)** Data represent mean ± SEM from three independent experiments performed in duplicate, multiple Mann-Whitney test, * p < 0.05, ** p < 0.01, *** p < 0.005 (relative to pcDNA3 condition). ƒ pvalue < 0.05, ƒƒ p < 0.01 and ƒƒƒ p < 0.005 (relative to anti-IFNAR condition).

## DISCUSSION

In this study, we identify CTIP2 as a key regulator of TLR3-driven antiviral responses in microglia. It is previously showed in the literature that TLR3 activation remove HIV latency in microglial cells ^46^. CTIP2 act via repression of NEAT1 and type I interferons (IFNs) to establish a negative feedback loop that links HIV-1 latency to innate immune control. Specifically, HIV-1 or poly(I:C) stimulation rapidly induces NEAT1 and IFN transcripts, which peak at 48 h, while CTIP2 expression rises later, reaching maximal levels at 72 h. Through IFNAR signalling, type I interferons themselves further sustain CTIP2 expression, which in turn represses NEAT1 and interferon production, thereby closing the loop. Our finding that CTIP2 represses NEAT1 and thereby dampens IFN output positions CTIP2 as a novel negative regulator of lncRNA-driven antiviral amplification. This is conceptually consistent with and extends previous work showing that lncRNAs can serve as either positive or negative modulators of IFN signalling. Indeed, recent reviews have highlighted that fine-tuning of IFN responses occurs by multi-layered control at the DNA, RNA, and protein levels, often via lncRNAs acting in feedback loops ^44^. NEAT1 is among the best-studied lncRNAs in this context. Its induction following viral infection or IFN stimulation has been documented in multiple systems, including influenza, HTNV, HIV, and herpesviruses ^42^. In many settings, upregulated NEAT1 supports the formation of paraspeckles, which act as subnuclear condensates that can sequester repressor proteins (e.g. SFPQ) or modulate promoter occupancy, thus shifting the balance of transcriptional regulation ^41^. For example, in a model of herpes simplex virus infection, NEAT1 was shown to influence viral gene transcription via chromatin and RNA-binding protein recruitment ^40^. In other settings, overexpression of NEAT1 increases paraspeckle formation, which can “buffer” or modulate the transcription of antiviral genes by sequestering factors like SFPQ away from promoters ^30^. Our findings go a step further, we show that CTIP2 not only modulates NEAT1 transcriptionally, but also physically inserts into paraspeckle RNPs and influences the assembly/stability of paraspeckle-associated complexes. Thus, CTIP2 bridges chromatin-level repression and RNA condensate-level organization. Notably, the dynamics we observe with NEAT1/IFN induction peaking earlier, followed by delayed accumulation of CTIP2, mirror the “aggregate-and-dissolve” cycles described in macrophages, where condensates form transiently to buffer stress and are then dissolved to reset transcriptional programs ^14^. Thus, CTIP2 may function as a “dissolver” or terminator of the paraspeckle-driven amplification phase in antiviral responses. As previously demonstrated, the HEXIM1-NEAT1-DNA-PK complex (termed HDP-RNP) has been shown to regulate IFN gene transcription after DNA sensing, where dissociation of paraspeckle components can free them to engage DNA-sensing pathways ^21, 41^. We propose that CTIP2 may favor or stabilize a paraspeckle-integrated repressive state. This implies another sequestration process, as CTIP2 depletion of NEAT1 coincides with relocation of SFPQ and related factors into CTIP2-induced nuclear structures, thereby reinforcing sequestration of activators and limiting further transcription. The delayed CTIP2 induction is central to our model, it allows an unrestrained early IFN wave (necessary for antiviral defense), followed by a controlled “braking” phase. The fact that CTIP2 is itself induced by type I IFN via IFNAR signalling closes a self-limiting loop that helps avoid excessive inflammation or autoimmune like overactivation. Importantly, we experimentally showed that blocking IFNAR1 attenuates CTIP2 induction and exacerbates NEAT1/IFN transcripts, supporting this negative feedback. One caveat is that while we see CTIP2 promoter occupancy (via ChIP-qPCR) of NEAT1 and recruitment of silencing factors (HDAC1 and HEXIM1/P-TEFb,), we have not yet mapped whether these factors are dynamically redistributed between the HIV LTR and the NEAT1 locus under stimulation. It remains possible that CTIP2 is not fully “evicted” from HIV chromatin but rather dynamically partitioned. Our glycerol gradient data suggest CTIP2 is required for the incorporation of paraspeckle proteins into heavier RNPs after cellular sensing of viral infection. However, paraspeckles are dynamic and heterogeneous in composition, CTIP2 may favor the formation of “repressive” paraspeckle-component containing structures resulting from the depletion of NEAT1 and the relocation of the proteins into CTIP2-induced structures. Additional high-resolution microscopy and proteomic profiling of CTIP2-associated RNPs may help deconvolve these subpopulations. Our findings, although derived primarily from a microglial cell line model, are consistent with our previous work showing that the effects of CTIP2 on transcriptional silencing and immune regulation are reproducible in other myeloid systems, including primary macrophages. This reinforces the notion that CTIP2 acts as a central regulator of viral latency across diverse myeloid reservoirs. Microglial cells, due to their longevity, self-renewal capacity, and anatomical isolation by the blood-brain barrier, represent one of the most stable HIV sanctuaries in the central nervous system ^5^. Similarly, tissue-resident macrophages contribute to long-term viral persistence and low-level reactivation events that sustain systemic inflammation despite effective antiretroviral therapy ^45^. In this context, understanding how CTIP2 orchestrates transcriptional repression and innate immune dampening is critical for strategies aimed at controlling viral latency and preventing reactivation within the CNS and peripheral reservoirs. Our study therefore strengthens the view that targeting CTIP2-regulated networks could be pivotal for durable HIV remission through simultaneous control of immune activation and maintenance of proviral silencing.

### Star Methods

#### Cell Culture, cell transfection and plasmids

HEK293T and human microglial cells (CHME5) were cultured in Dulbecco’s Modified Eagle Medium (DMEM; Gibco) supplemented with 10% fetal bovine serum (FBS, Gibco), 100 U/mL penicillin, and 100 µg/mL streptomycin at 37 °C in a humidified incubator with 5% CO□. For transient transfections, cells were seeded at 60-70% confluency and transfected with the indicated plasmids using JetPEI (PolyPlus) or JetPRIME (PolyPlus) reagents according to the manufacturer’s recommendations. Cells were maintained under standard culture conditions (37 °C, 5% CO□) for 24, 48, 72 or 96 h prior to downstream assays. The plasmids used in this study included pcDNA3 (empty control vector), full-length human FLAG-CTIP2, sh-CTIP2, and sh-Control constructs, which have been described previously^1,2^.

### Nuclear Protein Extraction

Nuclear protein extracts were prepared using successive osmotic shock with Buffer A and Buffer C. Briefly, cells were harvested from 10 cm dishes by scraping or trypsinization, pelleted at 1,200 rpm for 5 min at 4 °C, and washed twice with ice-cold PBS. Pellets were resuspended in 1 mL of ice-cold Buffer A (10 mM HEPES pH 7.9, 10 mM KCl, 1.5 mM MgCl□, 0.5 mM DTT) supplemented with protease inhibitor (Roche, 1:50), incubated on ice for 10 min, vortexed, and centrifuged at maximum speed of 1200 rpm for 1 min at 4 °C. Pellets were resuspended in 300 µL Buffer C (20 mM HEPES pH 7.9, 420 mM NaCl, 1.5 mM MgCl□, 0.2 mM EDTA, 0.5 mM DTT, 25% glycerol) supplemented with protease inhibitors, incubated on ice for 30 min, vortexed, and centrifuged at maximum speed 1200 rpm for 2 min at 4 °C. The supernatant was collected as nuclear protein extracts.

### Whole Cell Protein Extraction

Whole-cell lysates were prepared by resuspending washed cell pellets in 1 mL of ice-cold RIPA buffer (10 mM Tris-HCl pH 7.4, 150 mM NaCl, 1% Triton X-100, 0.1% SDS, 0.5% sodium deoxycholate, 1 mM EDTA), freshly supplemented with protease inhibitors (Roche, 1:50). Samples were incubated on a rotating wheel at 4 °C for 1 h, vortexed, and centrifuged at 10,000 rpm for 15 min at 4 °C. Supernatants were collected and stored at -20 °C.

Protein concentrations were determined using the Pierce™ BCA Protein Assay Kit (Thermo Scientific) following the manufacturer’s protocol. Briefly, samples and blanks were prepared in duplicates in 96-well plates (20 µL each), and 200 µL of reagent mix (50:1 ratio of Reagent A: Reagent B) was added. Plates were incubated at 37 °C for 30 min, cooled to room temperature, and absorbance was measured at 560 nm. Concentrations were calculated using a standard curve generated from bovine serum albumin standards.

### RNA Extraction and DNase Treatment

Total RNA was extracted using the NucleoSpin® RNA Plus kit (Macherey-Nagel) according to the manufacturer’s protocol with mechanical disruption and on-column genomic DNA removal. RNA binding and washing were performed with the provided buffers (WB1 and WB2), and RNA was eluted in 30 µL RNase-free water. Concentration and purity were assessed by NanoDrop spectrophotometry (A260/A280 ratio 1.9-2.1), and integrity was verified by agarose gel electrophoresis. Residual genomic DNA was removed using TURBO DNase (Invitrogen). RNA was incubated with DNase at 37 °C for 30 min, followed by heat inactivation at 75 °C for 10 min in the presence of 0.5 M EDTA. DNase-treated RNA was stored at -80 °C until further use.

### Cross-linking immunoprecipitation sequencing (CLIP-Seq)

This method has been described previously ^2^. Briefly, HEK293 and microglial cells were transfected with pFLAG-BCL11b or empty pcDNA3 control. The following day, cells were UV-crosslinked on ice (4000 J for 10 min) to stabilize RNA-protein interactions. Cells were lysed in buffer (50 mM Tris-HCl pH 7.4, 100 mM KCl, 2 mM MgCl□, 0.08 mM CaCl□, 0.1% SDS, 1% NP-40, 5 g/L deoxycholate) supplemented with RNase inhibitor (40 U/mL) and protease inhibitor cocktail. Lysates were treated with TURBO-DNase (10 U, 37 °C, 5 min) and centrifuged (13,000 rpm, 15 min, 4 °C). Supernatants were precleared sequentially with control beads and IgG-conjugated beads before overnight immunoprecipitation with anti-FLAG, anti-BCL11b, or IgG control antibodies at 4 °C. Beads were washed three times in high-salt buffer (50 mM Tris-HCl pH 7.4, 1 mM EDTA, 1 M KCl, 0.1% SDS, 1% NP-40, 5 g/L deoxycholate), followed by proteinase K digestion (2 mg/mL, 20 min, 37 °C). RNA-protein complexes were extracted with phenol/chloroform, and RNA was precipitated with sodium acetate (3 M, pH 5.5), Glycoblue, and 1:1 ethanol: isopropanol at -20 °C overnight. Precipitated RNA was pelleted by centrifugation (13,000 rpm, 10 min, 4 °C), washed twice with 80% ethanol, air-dried, and dissolved in 10 μL DEPC-treated water for sequencing.

### RNA Immunoprecipitation (RIP)

As previously described ^2^, microglial cells were cultured in 10 cm dishes to 80-90% confluence and lysed. Total RNA was extracted using the NucleoBond Xtra Maxi kit (Macherey-Nagel) following the manufacturer’s instructions. Lysates were cleared by centrifugation (10,000 rpm, 10 min, 4 °C) and precleared for 1 h at 4 °C with 25 μL Dynabeads Protein G (Invitrogen) bound to nonspecific IgG. For immunoprecipitation, 30 μL Dynabeads coated with 5 μg of the antibody of interest or control IgG were incubated with lysates overnight at 4 °C. RNA-protein complexes were recovered directly from beads by chloroform extraction, followed by centrifugation (13,000 rpm, 15 min, 4 °C). The aqueous phase was mixed with isopropanol and Glycoblue, incubated on ice (10 min), and pelleted by centrifugation (13,000 rpm, 10 min, 4 °C). Pellets were washed with 80% ethanol, air-dried, and resuspended in 16 μL DEPC-treated water. RNA was reverse transcribed using iScript Reverse Transcription Supermix (Bio-Rad) in a final 20 μL reaction. cDNA was diluted 1:5, and 2 μL was used per qPCR with SsoAdvanced Universal SYBR Green Supermix (Bio-Rad).

### Immunoprecipitation (IP)

For immunoprecipitation, nuclear or whole-cell extracts (1-1.5 mg protein) were incubated overnight with 4-5 µg of the indicated antibody or control IgG, coupled to 20 µL Dynabeads Protein G (Invitrogen). Beads were washed sequentially with low-salt buffer (IPLS: 50 mM Tris-HCl pH 7.5, 120 mM NaCl, 0.5 mM EDTA, 0.5% NP-40, 10% glycerol), high-salt buffer (IPHS: 50 mM Tris-HCl pH 7.5, 0.5% NP-40, 0.5 mM EDTA, 10% glycerol), and finally with IPLS. All buffers contained EDTA-free protease inhibitors (Roche). Bead-bound proteins were eluted by boiling in SDS sample buffer and analysed by Western blotting.

### Mass Spectrometry (LC-MS/MS)

As previously described ^2^, clear protein extracts were subjected to anti-FLAG immunoprecipitation. Immunoprecipitated proteins were eluted, digested with trypsin, and processed. Peptides were separated by reverse-phase liquid chromatography using an Ultimate Ultra3000 chromatography system (Thermo Scientific, Germany) and analyzed on a high-resolution Q-Exactive mass spectrometer (Thermo Scientific). Raw spectra were processed with MaxQuant software, and protein intensities were calculated by summing peptide ion intensities uniquely corresponding to each protein sequence across all samples. Interacting proteins were determined after normalization to trypsin digestion controls and to an empty vector (pcDNA3) transfection control. The data represent three independent experiments performed in duplicate.

### Western Blotting

Protein samples (20-40 µg) were denatured in 4× Laemmli buffer with 10% β-mercaptoethanol, heated at 95 °C for 5 min, and resolved on 8-10% polyacrylamide gels (Bio-Rad Mini-PROTEAN). Proteins were transferred onto nitrocellulose membranes using the Bio-Rad Trans-Blot Turbo Transfer system. Membranes were blocked in EveryBlot Blocking Buffer (Bio-Rad) for 5 min, incubated with primary antibodies (**Table S1**) for 1 h at room temperature, washed in Tween-PBS (5 times for 5 minutes each on a shaking platform), and incubated with HRP-conjugated secondary antibodies for 1 h. Signals were detected with SuperSignal Chemiluminescence (Thermo Fisher) and visualized using the Fusion imaging system (Vilber Lourmat).

### Glycerol Gradient Sedimentation Analysis

This method has been described before ^3^. We followed it with some minor changes. Nuclear extracts (250 µL, 1.5 mg protein) were layered onto discontinuous 10-40% glycerol gradients (4.5 mL) and centrifuged for 5 h at 32,000 rpm, 4 °C (SureSpin 632 rotor, Thermo Scientific). Twenty-three fractions (200 µL each) were collected from the top and analyzed by SDS-PAGE and Western blot. Sedimentation buffer contained 10 mM Tris-HCl pH 8.0, 150 mM NaCl, 10 mM KCl, 1.5 mM MgCl□, 0.5% Triton X-100, 0.5 mM EDTA, and 1 mM DTT, freshly supplemented with Roche Complete protease inhibitors.

### Chromatin-IP (ChIP)

ChIP was performed to assess protein-NEAT1 promoter interactions. Briefly, cells were crosslinked with 1% formaldehyde for 10 min at RT, quenched with 125 mM glycine, washed, and collected in cold PBS. Nuclei were isolated by lysis in buffer containing protease inhibitors, and chromatin was fragmented (∼200bp) with MNase (200U per million of cells, M0247SVIAL, New England Biolabs) ∼200. After clarification by centrifugation, chromatin was incubated overnight at 4 °C with protein A/G magnetic beads pre-bound to specific antibodies or IgG controls (**Table S1**). Immune complexes were sequentially washed in low-salt, high-salt, LiCl, and TE buffers, then eluted in SDS buffer. Crosslinks were reversed by heating at 65 °C for 6 h, followed by RNase A and proteinase K digestion. DNA was purified using phenol-chloroform extraction and analyzed by quantitative PCR with primers targeting internal (P1) and external (P0) regions of NEAT1 (**Table S2**) as described in ^39^. Enrichment was calculated relative to input chromatin.

### Reverse Transcription-quantitative PCR (RT-qPCR)

RTqPCR was used to measure transcript levels. Total RNA was isolated using a silica-membrane spin column kit (NucleoSpin RNA, Macherey-Nagel, # 740955) according to the manufacturer’s protocol, including on-column DNase I digestion to remove genomic DNA. RNA concentration and purity were assessed spectrophotometrically (A260/A280 ratio). Between 500 ng of RNA was reverse transcribed using random hexamers and oligo(dT) with a commercially available reverse transcriptase (iScript cDNA synthesis kit, Bio-rad, #1708890) in a 20 µL reaction. qPCR was performed on a real-time instrument (CFX96, Bio-rad) using SYBR Green master mix in 20 µL reactions with gene-specific primers (primer sequences listed in supplementary **Table S2**). Thermal cycling: 95 °C for 2-3 min, then 40 cycles of 95 °C for 10-15 s and 60 °C for 30-60 s; include a melt-curve. Relative quantification used the ΔΔCt method with GAPDH as housekeeping controls; all reactions were run in technical triplicates and included no-template and no-RT controls. Data are presented as mean ± SEM from at least four biological replicates.

### Statistical analysis

For all datasets, the Gaussian distribution was assessed using the Shapiro-Wilk normality test. If the data followed a Gaussian distribution, statistical differences were evaluated using an unpaired t-test (with Welch’s correction for unequal variances). For non-Gaussian distributions, the Mann-Whitney test or Friedman test were employed to determine statistical significance. All analyses were conducted using GraphPad Prism software (version 9.1.1), and results are expressed as mean ± SEM. Statistical significance was set at p < 0.05, with levels denoted as *p < 0.05, **p < 0.01, ***p < 0.001, ****p< 0.0001.

Antibodies List.

Antibodies List.
| S.No | Antibody | Company Name | Reference No. |
| --- | --- | --- | --- |
| 1 | Flag | Sigma | F3165 |
| 2 | Anti-Bcl11b | Bethyl | A300-384A |
| 3 | PSP (G-7) SFPQ | SANTA CRUZ | sc-37502 |
| 4 | P54/nrb (NONO) | SANTA CRUZ | sc-166702 |
| 5 | PSPC1 | SANTA CRUZ | sc-374181 |
| 6 | FUS/TLS | SANTA CRUZ | sc-47711 |
| 7 | HEXIM1 (D-8) | SANTA CRUZ | sc-390059 |
| 8 | CDK9 (D-7) | SANTA CRUZ | sc-13130 |
| 9 | KLF16 (F-4) | SANTA CRUZ | sc-377519 |
| 10 | Actin | SANTA CRUZ | sc-58673 |
| 11 | Anti-Mouse | SANTA CRUZ | sc-516102 |
| 12 | Anti-Rabbit | Thermo Scientific | 32460 |
| 13 | IRF3 | Invitrogen | MA5-32348 |
| 14 | Phospho-IRF3 | Invitrogen | PA5-36775 |
| 15 | Cyclin T1 | Bethyl | A303-498A |
| 16 | P65 | Abcam | Ab7970-1 |
| 17 | Sp-1 | Millipore | LOT:3921951 |
| 18 | HDAC1 | Millipore | LOT:4243306 |

**Table S2:**
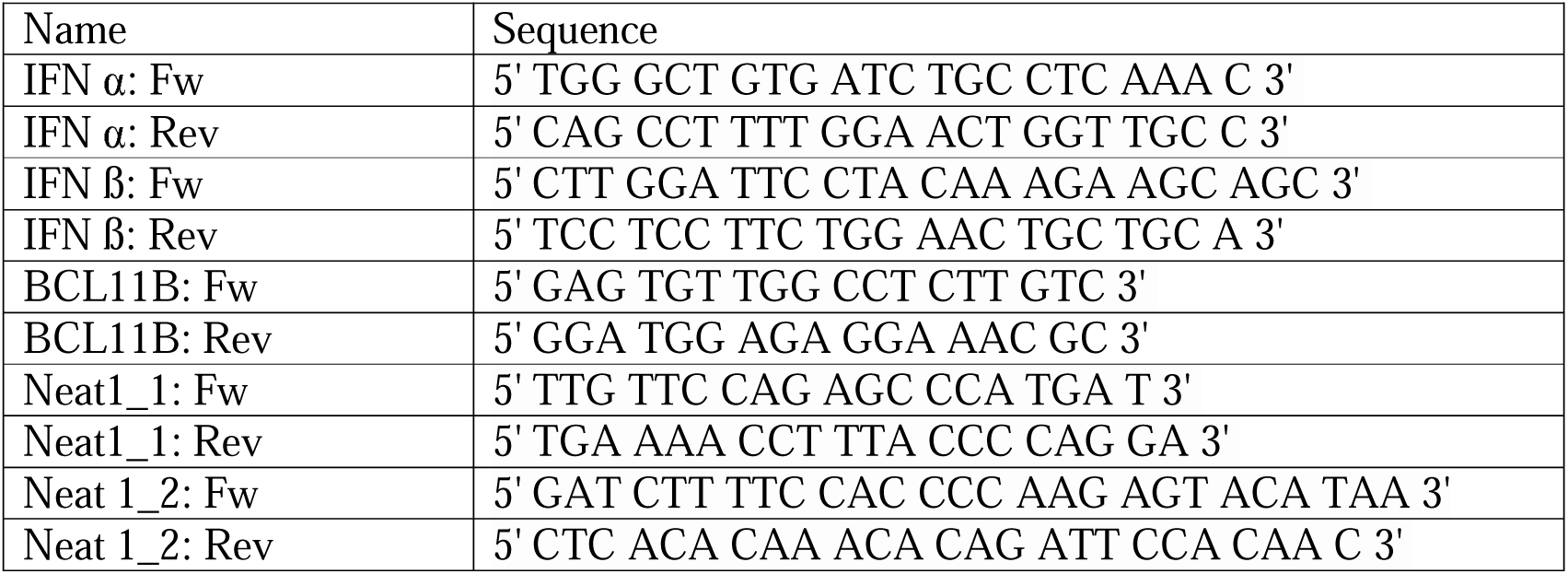
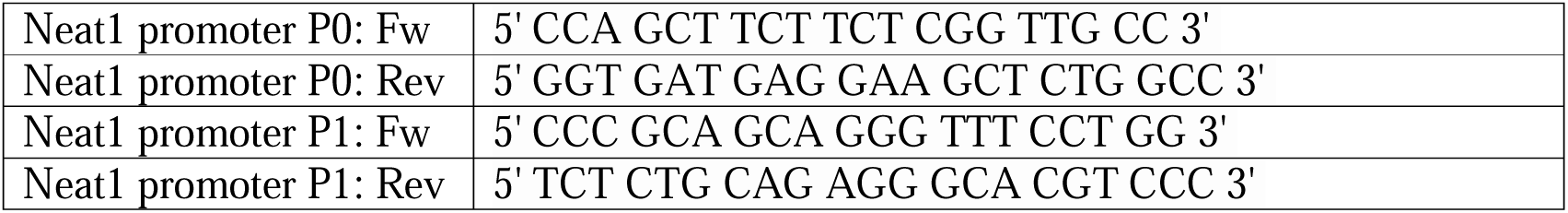
Primers List.

