## Supplementary data for "THE HIV-1 SILENCER BCL11B/CTIP2 REGULATES THE TLR3-MEDIATED CELLULAR RESPONSE TO VIRAL INFECTIONS"

**Supplementary Figure 1**

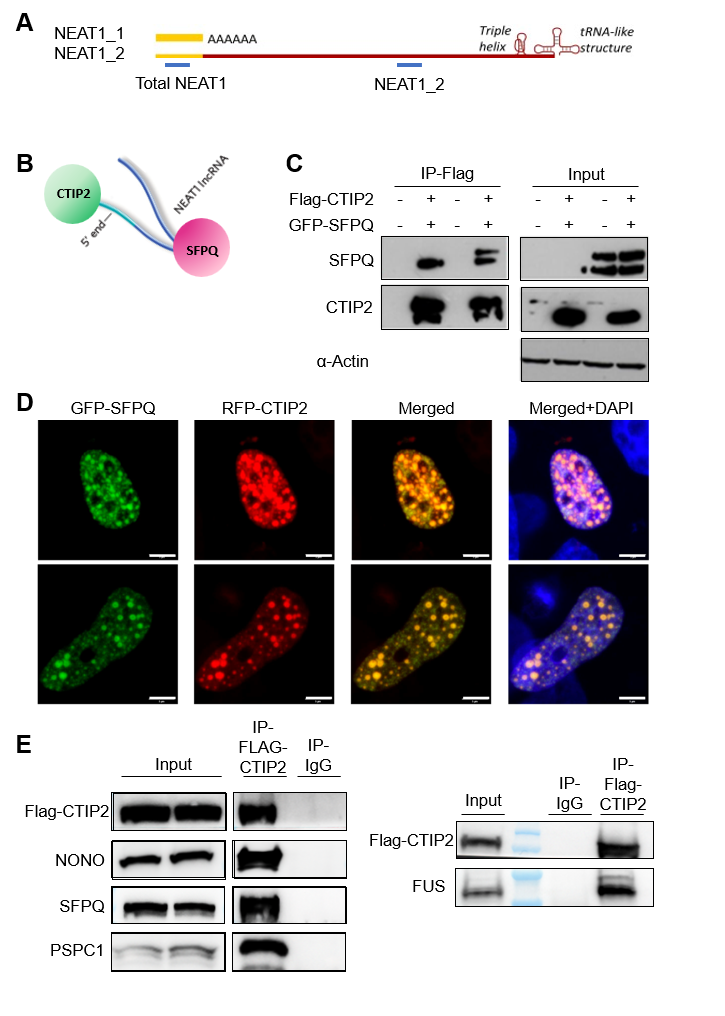

**Supplementary Figure 2**

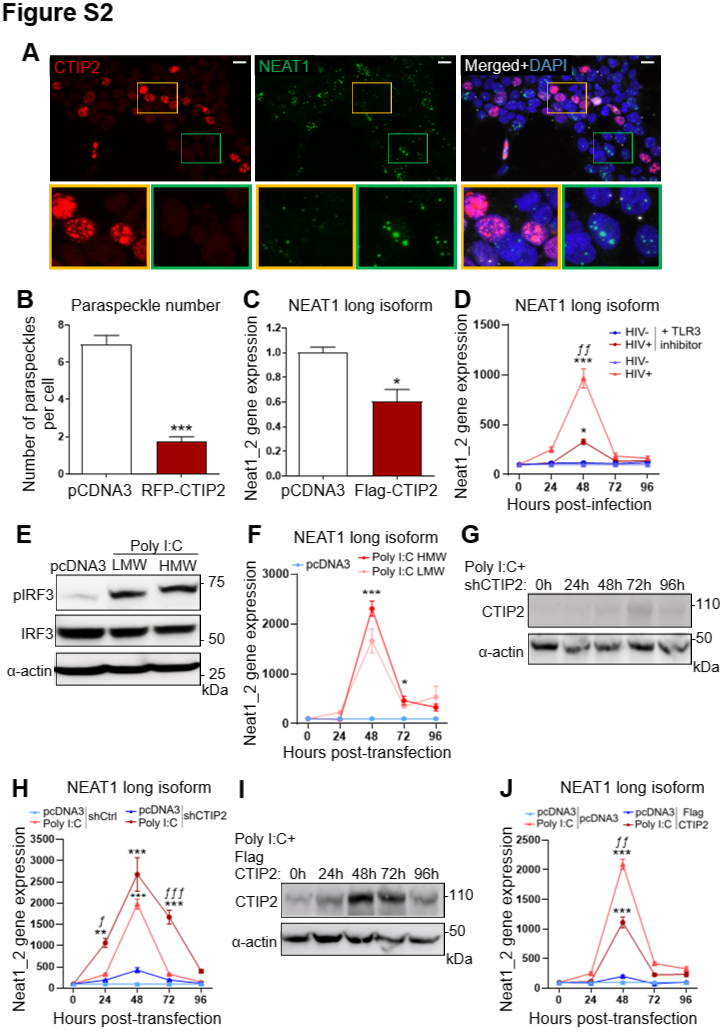

**Supplementary Figure 3**

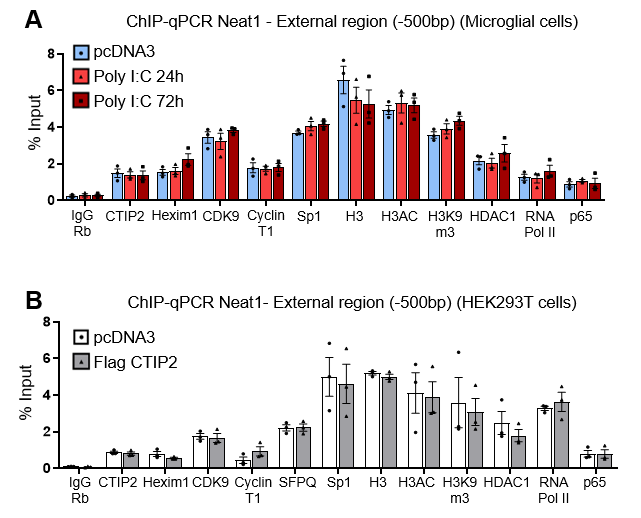

**Supplementary Figure** **1**

**Figure S1. (A)** Schematic of NEAT1 isoforms. NEAT1_1 terminates with a canonical poly(A) tail (yellow), whereas NEAT1_2 extends further and contains 3′-end structural motifs, including a triple helix and tRNA-like elements (red). **(B)** Domain-specific binding preferences of CTIP2 and SFPQ along the NEAT1 transcript. CLIP-seq and RIP-qPCR analyses show CTIP2 enrichment at the 5′ region, while SFPQ preferentially associates with central and 3′ domains of NEAT1_2. **(C)** Co-immunoprecipitation of CTIP2 with SFPQ. HEK293 cell lysates transfected with Flag-CTIP2 were subjected to immunoprecipitation with anti-Flag antibody or control IgG and analyzed by Western blotting, confirming the association with SFPQ. **(D)** Confocal microscopy images shows HEK293 cells expressing RFP-tagged CTIP2 and SFPQ (green) localized to nuclear foci, with colocalization of CTIP2 (red) within paraspeckle structures. Nuclei were counterstained with DAPI (blue). **(E)** Co-immunoprecipitation of Flag-CTIP2 with paraspeckle proteins. Immunoprecipitation from HEK293 cells confirmed stable association of CTIP2 with NONO, SFPQ, PSPC1, and FUS.

**Supplementary Figure** **2**

**Figure S2. (A)** RNA FISH and IF staining respectively for NEAT1 (green) and CTIP2 in microglial cells transfected with CTIP2-RFP or mock transfected (pcDNA3). **(B)** Quantification of RNA FISH NEAT1-positive nuclear foci in microglial cells transfected with CTIP2-RFP or pcDNA3. Quantification across 50 cells per condition.

**(C)** qRT-PCR analysis of total NEAT1 transcripts in microglial cells transfected with Flag-CTIP2, showing reduced expression compared to pcDNA3 controls. (B-C) Data represent mean ± SEM; Student’s t-test (***p < 0.001; *p < 0.05). **(D)** qRT-PCR of NEAT1_2 expression in HIV-1 infected (HIV+) or mock-treated (HIV-) microglial cells, in the presence or absence of a TLR3/dsRNA inhibitor (100 µM). HIV-1 induced strong NEAT1_2 upregulation, peaking at 48 h, which was abolished by TLR3 inhibition. **(E)** Western blot analysis of IRF3 activation in microglial cells transfected with pcDNA3 or poly(I:C). Poly(I:C) induced robust phosphorylation of IRF3 (p-IRF3), without altering total IRF3 levels, consistent with post-translational activation downstream of TLR3 signaling. α-actin served as a loading control. **(F)** qRT-PCR of NEAT1_2 transcripts in microglial cells transfected with HMW or LMW poly(I:C), or pcDNA3 as control. Both duplexes induced NEAT1_2, with stronger induction by HMW poly(I:C). No induction was detected in controls. **(G)** Western blot validation of CTIP2 knockdown in microglial cells co-transfected with poly(I:C) and shCTIP2, confirming loss of CTIP2 protein. α-actin served as a loading control.

**(H)** qRT-PCR of NEAT1_2 in microglial cells co-transfected with poly(I:C) and shCtrl or shCTIP2. CTIP2 knockdown significantly enhanced and prolonged NEAT1_2 induction compared to shCtrl. **(I)** Western blot validation of CTIP2 overexpression in microglial cells transfected with poly(I:C) and Flag-CTIP2, showing strong CTIP2 protein accumulation at 48 (Flag-CTIP2) and 72 h (endogenous CTIP2). α-actin served as a loading control.

(J) qRT-PCR of NEAT1_2 transcripts in microglial cells transfected with poly(I:C) and either pcDNA3 or Flag-CTIP2. While poly(I:C) induced robust NEAT1_2 expression in control cells, overexpression of Flag-CTIP2 significantly repressed this induction.

(D, F, H, J) Mean ± SEM, n=4 independent experiments per time point and per conditions, multiple Mann-Whitney test. * p < 0.05, ** p < 0.01, *** p < 0.005, **** p < 0.001 (relative to pcDNA3 condition). ƒ pvalue < 0.05, ƒƒ p < 0.01 and ƒƒƒ p < 0.005 (relative to TLR3 inhibitor (D), shCtrl (H) and pcDNA3 (J) conditions).

**Supplementary Figure** **3**

**Figure S3 legend. (A)** ChIP-qPCR analysis of microglial cells transfected with pcDNA3 or stimulated with poly(I:C) (1 µg/µL) for 24 h or 48 h. Occupancy of indicated proteins Xbp 5’ before the NEAT1 promoter region was quantified. Data represent three independent experiments performed in duplicate and are shown as percentage of input. Mean ± SEM, Mann–Whitney tests (*p ≤ 0.05; **p ≤ 0.01). **(B)** ChIP-qPCR analysis of HEK293T cells transfected with pcDNA3 or Flag-CTIP2 for 48 h. Binding of indicated proteins to the NEAT1 promoter was quantified. Data represent three independent experiments performed in duplicate and are shown as percentage of input. Mean ± SEM, Mann–Whitney tests (*p ≤ 0.05; **p ≤ 0.01; ***p ≤ 0.001).
